# Automated EEG mega-analysis II: Cognitive aspects of event related features

**DOI:** 10.1101/411371

**Authors:** Nima Bigdely-Shamlo, Jonathan Touryan, Alejandro Ojeda, Christian Kothe, Tim Mullen, Kay Robbins

## Abstract

In this paper, we present the results of a large-scale analysis of event-related responses based on raw EEG data from 17 studies performed at six experimental sites associated with four different institutions. The analysis corpus represents 1,155 recordings containing approximately 7.8 million event instances acquired under several different experimental paradigms. Such large-scale analysis is predicated on consistent data organization and event annotation as well as an effective automated pre-processing pipeline to transform raw EEG into a form suitable for comparative analysis. A key component of this analysis is the annotation of study-specific event codes using a common vocabulary to describe relevant event features. We demonstrate that Hierarchical Event Descriptors (*HED* tags) capture statistically significant cognitive aspects of EEG events common across multiple recordings, subjects, studies, paradigms, headset configurations, and experimental sites. We use representational similarity analysis (*RSA*) to show that EEG responses annotated with the same cognitive aspect are significantly more similar than those that do not share that cognitive aspect. These *RSA* similarity results are supported by visualizations that exploit the non-linear similarities of these associations. We apply temporal overlap regression to reduce confounds caused by adjacent events instances and extract time and time-frequency EEG features (regressed ERPs and ERSPs) that are comparable across studies and replicate findings from prior, individual studies. Likewise, we use second-level linear regression to separate effects of different cognitive aspects on these features, across all studies. This work demonstrates that EEG mega-analysis (pooling of raw data across studies) can enable investigations of brain dynamics in a more generalized fashion than single studies afford. A companion paper complements this event-based analysis by addressing commonality of the time and frequency statistical properties of EEG across studies at the channel and dipole level.

## 1. Introduction

The main goal of this work is to establish the feasibility of large-scale EEG (electroencephalography) mega-analysis (pooling of raw data across studies) by jointly analyzing multiple EEG studies representing different paradigms, headsets, subjects, and experimental sites. This work is motivated by the success of data-pooling efforts for other neuroimaging modalities such as fMRI (functional magnetic resonance imaging) (Costafreda, 2009) and the need to establish reproducible, generalizable observations in EEG (Jas et al., 2018).

EEG as a neuroimaging modality provides some unique capabilities such as the ability to record behavior in natural settings. However, registration of EEG measurements is complicated by the significant variability introduced by subject motion, muscle activity, eye activity, headset placement, signal transmission media (hair, scalp, and skull), as well as recording issues such as loose or damaged sensors. Intrinsic differences in the recording systems, themselves, as demonstrated by Melnik et al. (Melnik et al., 2017) also introduce significant variability. Compared to fMRI, there is less agreement on recording standards for preprocessing of EEG data or on methods of reporting results, making it difficult to perform meta-analyses except at very qualitative levels.

The lack of agreed-upon standards for EEG suggests that pooling of raw EEG data across studies may be more promising than meta-analysis (pooling of analysis results) because it eliminates the significant variability introduced by differing choices in preprocessing steps (e.g., referencing, filtering, artifact removal). This paper adopts a particular data organization and an automated preprocessing suite to facilitate computation (Bigdely-Shamlo et al., 2015) (Bigdely-Shamlo et al., 2016a) (Bigdely-Shamlo et al., 2016b) (Kleifges et al., 2017). However, the main results of this paper are not strictly tied to the preprocessing methodology. Rather, the important element is the uniformity of data processing in a “reasonable” but automated (i.e., identical) manner. Based on the assumption that “clean” EEG can be produced in some standardized fashion, we consider three important questions regarding mega-analysis:

1. How can the common properties of events be isolated and compared across experiments that differ in many details?
2. How can the effects of adjacent events be isolated in the face of overlapping responses and potential confounds within individual experiments?
3. How can variability introduced by subject, headset, and other factors be properly identified and partitioned to understand the fundamental nature of the underlying neural responses.

We investigate the first question by introducing the notion of a “cognitive aspect”, which is specified by annotating experiment-specific event codes using a common ontology called Hierarchical Event Descriptors (*HED* tags). We then analyze these cognitive aspects across studies to isolate particular effects based on concepts rather than event codes. To address the second question, we use an implementation of temporal overlap regression similar to that proposed by Kristensen et al. (Kristensen et al., 2017) with some enhancements to support large-scale automated analysis. We address the third issue by augmenting this regression with a hierarchical organizational model. The model allows separation of subject, headset, paradigm, and cognitive aspects in a manner that is scalable and generalizable across diverse collections of EEG datasets. The paper provides several demonstrations of the efficacy of this approach in comparison with traditional analyses. The paper also uses representational similarity analysis (*RSA*) (Kriegeskorte et al., 2008) to assess the statistical significance of the relationship between cognitive aspects and EEG signal patterns, as well as t-distributed stochastic neighbor embedding (*t-SNE*) (Van Der Maaten and Hinton, 2008) to visualize these relationships.

The paper is organized as follows. The Methods section describes the experimental data and introduces the notion of cognitive aspects as realized by *HED* tags. After briefly describing the automated pipeline that facilitates processing, the paper describes the implementation of temporal overlap regression and second-level hierarchical modeling. The paper also defines the EEG signal pattern and *HED* tag presence dissimilarity measures needed to apply *RSA*. The Results section follows a similar organization with a presentation of results on the information content and statistical properties of cognitive aspects across heterogeneous studies. We then provide a comparison of traditional averaging versus overlap regression in the context of multiple studies and use a second-level linear model to extract patterns associated with *HED* tags that are common across studies. Finally, we use *RSA* to show which *HED* tags are associated with EEG signal patterns that are repeated consistently across studies. The Discussion section offers a perspective on how these results compare to previous reports from individual studies as well as directions for future research.

## 2. Methods

Table 1 summarizes the 17 studies used in this analysis. The appendix of a companion paper (Bigdely-Shamlo et al., 2018) provides additional details about these studies. All studies were conducted with voluntary, fully-informed subject consent and were approved by the Institutional Review Boards of the respective institutions. Each EEG dataset was converted to an EEGLAB EEG structure (Delorme and Makeig, 2004) (Delorme et al., 2011).

**Table 1.**
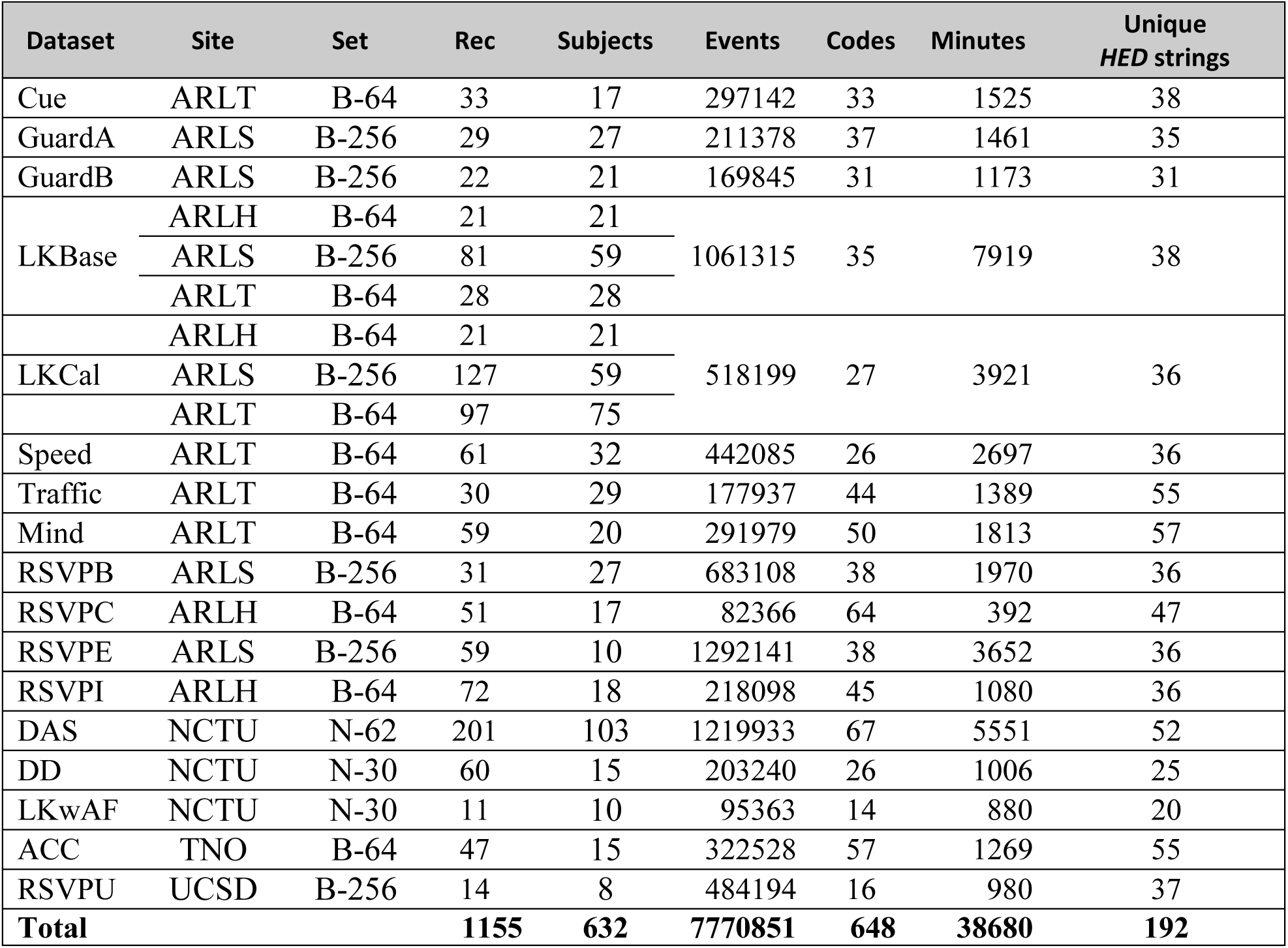
Summary of datasets used. Subjects may overlap across studies at the same site. The Set column (B = BioSemi, N = Neuroscan) specifies headset type and number of channels.

The event types in each study were identified by study-specific event codes provided by the original experimenters. The data from the studies was otherwise unprocessed, with one exception. The data acquired from Neuroscan headsets (N-30 and N-62) was automatically referenced to mastoids as part of the acquisition process.

### 2.1 Data organization and preprocessing

As mentioned in the Introduction, standardization of EEG for data sharing is less mature than for fMRI, and standards for preprocessed EEG have not yet been universally adopted. However, the two essential elements required for mega-analysis of the type proposed in this paper are the availability of data that has been effectively cleaned using the same automated pipeline and events that have been annotated using a common event annotation system. This subsection describes the particular automated pipeline we used to organize and preprocess the data, which is the starting point of the analysis. We describe the event annotation process in a subsequent subsection.

We used several open-source tools that we have developed to do the automated preprocessing. To facilitate preprocessing we organized each study by placing it in a containerized format using the EEG Study Schema (ESS) as described in Bigdely-Shamlo et al. (Bigdely-Shamlo et al., 2016b). ESS, which is part of the BigEEG technology stack (*BigEEG Workflow*, 2018), allows specification, in machine-readable format, of channel labels and locations as well as meta-data such as subject and task information about each recording.

We then applied the *PREP* pipeline to all studies (Bigdely-Shamlo et al., 2015). After removing line noise from each data recording, *PREP* interpolates noisy channels and then calculates and removes the robust average reference from each recording. Next, our pipeline removes non-EEG channels from each recording, assigns 10-20 labels to channels that do not have standard labels (e.g., 256-channel BioSemi caps) based on channel distances to standard 10-20 locations (Jurcak et al., 2007), and selects a maximum of 64 channels based on the 64-channel standard 10-20 configuration. We down-sampled to 128 Hz and applied a 1690-point high-pass filter at 1 Hz using the *pop_eegfiltnew()* function of EEGLAB to remove low-frequency drifts.

The next stage of preprocessing deals with eye events and eye artifacts. First, we apply *BLINKER* (Kleifges et al., 2017) to channel data to identify latencies associated with different phases of blinks and insert corresponding event markers into `EEG.event`. *BLINKER* also extracts a continuous “blink” signal that follows the blink-induced EEG. We used the inserted blink events and the continuous “blink” signal in later processing.

We used a combination of independent component analysis (ICA) and regression to remove eye movement activity. After performing Infomax ICA (Bell and Sejnowski, 1995) on data not containing outlier amplitudes as described in Mullen et al. (Mullen et al., 2015), we applied *EyeCatch* (Bigdely-Shamlo et al., 2013) to identify and remove independent components (ICs) associated with eye movements. We also regressed out both the effects of blink events and the continuous blink signal returned from *BLINKER*. This regression used the same temporal overlap procedure as described in the next section. However, blinks are assumed to be fixed patterns within an interval of [-1, 1] seconds time-locked to the peak amplitude of the blink, while the other event-related analyses presented in this paper used intervals of [-2, 2] seconds time-locked to the event onset. Except where otherwise noted, the analyses described in this paper used the following subset of 26 channels common across all of the recordings: Fp1, Fp2, F3, Fz, F4, F7, F8, FC3, FCz, FC4, FT7, FT8, C3, Cz, C4, TP7, TP8, CP3, CPz, CP4, P3, Pz, P4, O1, Oz, and O2.

### 2.2 Extraction of event-related features at the recording level

We extracted four types of event-related features from continuous EEG data as shown by the pipeline of Figure 1. Time domain features include traditional event related potentials computed by averaging (ERPs) and by temporal overlap regression (rERPs). Time-frequency domain features include event-related spectral perturbations computed by averaging (ERSPs) and by temporal overlap regression (rERSPs). We gathered these features for all the recordings into a single database (*LARG*) for efficient analysis and hierarchical modeling.

**Figure 1.**
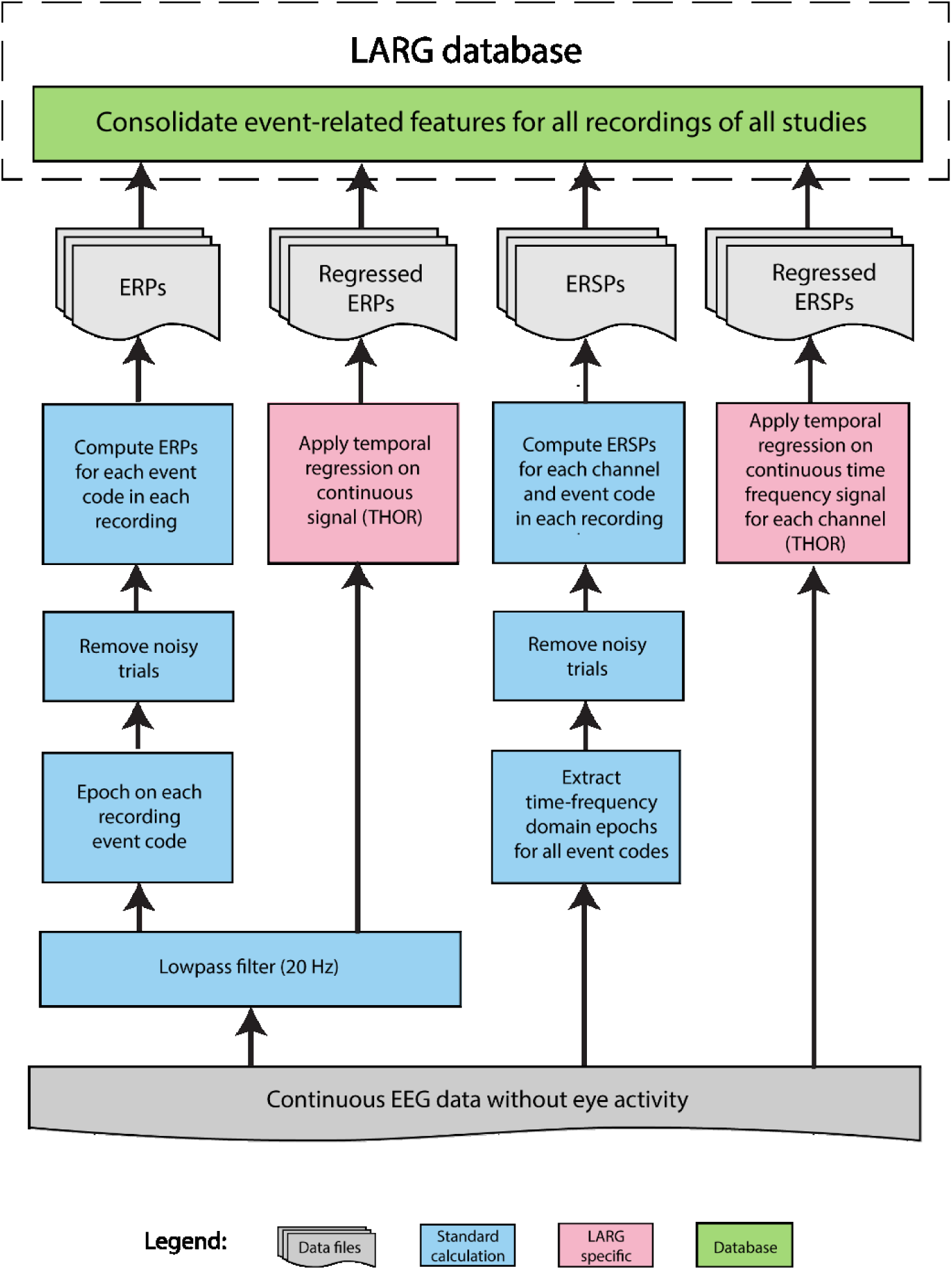
Overview of the pipeline for computing event-related features.

We computed event-related features based on intervals of [-2, 2] seconds time-locked to the specified events, either by averaging (for traditional ERPs and ERSPs) or by incorporation into a design matrix (for regression-based rERPs and rERSPs).

To compute ERPs, we first low-pass filtered the data at 20 Hz using the *pop_eegfiltnew()* function from EEGLAB and then extracted trials time-locked to each event code in each recording. We randomly selected a maximum of 1000 instances of each event code in each recording, with no two selected event instances corresponding to the same code allowed to be closer than 0.5 seconds. This sub-selection is crucial for processing data from Rapid Serial Visual Presentations (RSVP) experiments, which often include thousands of non-target events in close proximity to each other.

To detect outlier epochs, we concatenated all selected trials of a recording together to form a single *channels* × *time* array and robust z-scored each channel (subtracting the channel median and dividing by 1.4826 × the median absolute deviation for that channel). We then formed an epoch rejection mask by marking those epochs whose mean (over *channels* × *time*) absolute z-score was greater than 3 as outlier trials. We calculated ERPs by averaging unnormalized non-outlier trials for each event type in each recording. The z-score normalization was only used for outlier determination.

To generate time-frequency data, we applied the discretized continuous wavelet transform (using the MATLAB *cwt* function) to the continuous EEG time course associated with each channel to obtain a time-varying amplitude spectrogram (square root of power of 50 frequencies logarithmically sampled between 2 and 40 Hz). We scaled the resulting amplitudes by subtracting the median and then dividing by 1.4826 × the median, with median computed separately at each frequency over all time points for each recording.

We concatenated the time courses of the selected trials in a recording for each (*channel, frequency*) pair separately, performed a robust z-scoring across time separately for each (*channel, frequency*) pair, and marked trials with mean absolute robust z-score (over *time* × *frequency* × *channel*) greater than 3 as outlier trials. As with ERPs, z-score normalization is only used for outlier determination. We calculated ERSPs by averaging the unnormalized non-outlier trials for each event type and each channel in each recording. Because outlier detection requires all of the trials to be kept in memory simultaneously, we down-sampled the trials to 40 Hz for the ERSP calculations due to performance considerations.

We computed regression-based features (rERPs and rERSPs) on continuous data separately for each recording, excluding event types in a recording with fewer than 10 instances or events types whose instances frequently coincide with other events. For the analysis of this paper, we implemented a “BigEEG friendly” version of the regularized temporal overlap regression algorithm described in (Kristensen et al., 2017) for regularized temporal regression-based feature extraction. We refer to this algorithm as Temporal Hierarchical Overlap Regression (*THOR*). *THOR* includes an iterative method for finding and ignoring time points having outlier values. On each iteration, *THOR* fits a general linear model trained on non-outlier values identified in the previous iteration and computes regression residuals. It then marks a time point as an outlier (by removing it from the regression) when either (a) the robust z-score of the residual (Euclidean) amplitude is higher than 30, or (b) the smoothed (with a 120-point forward and backward box smoothing) residual amplitude has a robust z-score higher than 5. The iterations continue until either less than 0.001 of the time points change their outlier designation in an iteration or 50 iterations complete. While our pipeline takes continuous EEG with eye artifacts removed, muscle and other artifacts remain, and this iterative procedure allows *THOR* to reduce model errors due to artifacts.

*THOR* uses the Generalized Cross Validation (GCV) (Golub et al., 1979) (Chung and Español, 2017) method to select the regularization (or ridge) parameter, *λ*, for the ridge regression model. GCV can be viewed as a rotation-invariant version of the ordinary cross validation method (also known as leave-one-out cross-validation) that seeks to find the value of *λ* that, given a sample of *n* measurements, minimizes the mean square error of the prediction of each measurement from a model fit to the remaining *n* - 1 measurements. The optimization of the GCV curve yields a *λ* that minimizes the tradeoff between the prediction error and the complexity of the model, thereby guarding against overfitting. The GCV is a model selection criteria that has been extensively used in the brain imaging literature (Subramaniyam et al., 2010) (Hu et al., 2018).

We gather the resulting event-related patterns or features (ERPs, ERSPs, rERPs, rERSPs) into a database to facilitate analysis. Each feature is associated with a study-specific event code and a recording ID. These features also have as associated metadata, subject information and the *HED* tags associated with the event code. The *HED* tags describe the essential elements of events in a study-independent way as described in the next section.

### 2.3 Capturing cognitive aspects of events using *HED* tags

EEG studies typically use study-specific codes to annotate events. Processing a heterogeneous collection of datasets requires manual translation of codes into a common language. This process is more complex than just translating all potential “target” events to have an event code “target.” In one study, the target could be a red square in one block of trials and a blue triangle in another. Another study could use faces rather than shapes as targets, or auditory tones rather than visual stimuli. Are these differences important?

The *HED* (Hierarchical Event Descriptor) system (Bigdely-Shamlo et al., 2016a) proposes a semi-structured, standardized vocabulary for annotating events using comma-separated strings of *HED* tags (paths from the *HED* schema hierarchy) (*HED-schema*, 2018). A red triangle target event might have tags:

> *Participant/Effect/Cognitive/Target*
>
> *Item/2D shape/Triangle*
>
> *Attribute/Visual/Color/Red*

The *HED* event tagging system enables complex queries to look for patterns across studies – one could focus on all targets, or just on red targets, or on triangles. Paradigm-specific details can be annotated explicitly and then processed or ignored during analysis. Some analyses require the unrolling of *HED* tags into their component path prefixes. For example *Attribute/Visual/Color/Red* is unrolled into the following path prefixes: *Attribute/Visual/Color/Red, Attribute/Visual/Color, Attribute/Visual*, and *Attribute.* Each path prefix is itself a valid *HED* tag, encompassing a broader spectrum of event types.

The underlying assumption for *HED* tagging is that each event is associated with zero, one, or more cognitive phenomena, and these phenomena are relatively separable. We refer to these cognitive phenomena as *cognitive aspects*. Figure illustrates how cognitive aspects, as captured by *HED* tags, disassociate event properties from study-specific codes to facilitate the identification of cognitive effects across studies. The *Oddball* aspect captured by *HED Tag 1* only documents event code *A*_*1*_ of study A, while the *Error feedback* aspect captured by *HED Tag 2* documents event code *A*_*n*_ of Study A and event code *Z*_*1*_ of Study Z, but not event codes *A*_*1*_ or *Z*_*m*_ of those respective studies. The *Visual effect* aspect captured by *HED Tag q*, on the other hand, documents event *A*_*1*_ of Study A and both event codes *Z*_*1*_ and *Z*_*m*_ of Study Z. Thus, the *HED* tagging mechanism enables the use of automated methods to isolate cognitive aspects such as *Error feedback* across different events within and across studies. This disassociation is essential for studying complex effects on a large scale.

We use Shannon entropy to characterize the diversity of event instances associated with each tag, i.e. how many different event codes these encompass and in which proportions. The more diverse the events associated with a tag are, the less likely it is that confounds will emerge (due to idiosyncrasies of patterns associated with each event code). Let *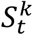* be the number of occurrences of *HED* tag *k* for a particular study-specific event code, *t*. Then is the probability 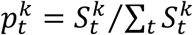 that an event instance matching the tag *k* is from an event with event code *t*. The Shannon entropy (in nats) is:

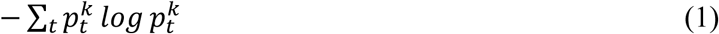

Here we have assumed that events with different codes are truly different. We also unroll each *HED* tag into its component path prefixes for the analysis, since the depth of a given *HED* tag is determined (somewhat arbitrarily) by the level of detail provided by the annotator.

The experimenters or their support staff initially annotated the study-specific event codes using the *HED* tag system. Since the quality of multi-study analysis depends on the consistency of the assignment of *HED* tags to study-specific event codes by annotators, two people who did not perform the initial annotation reviewed the annotations across all of the studies for consistency and made several corrections after discussion with the experimenters and annotators. In some cases the *HED* hierarchy itself (*HED-schema*, 2018) was modified to accommodate the needed vocabulary. Once a consensus mapping of study-specific event codes to *HED* tags was created, we included the annotations for the individual events in the `EEG.event.usertags` field of the EEGLAB EEG structure for each recording.

### 2.4 Common cognitive aspects of event-related features across studies

Each extracted event-related pattern (ERP, ERSP, rERP, rERSP) is a model of brain dynamics for a single study-specific event code and a particular recording. Likewise, each event code is associated with a set of *HED* tags as illustrated in Figure 2. A straightforward way to understand commonality and variability of these patterns and their relationship to underlying cognitive aspects across studies is to average patterns labeled with a particular *HED* tag. We call this analysis method aspect averaging (*AA*). Each tag is unrolled into its path prefixes to select the patterns to average.

**Figure 2.**
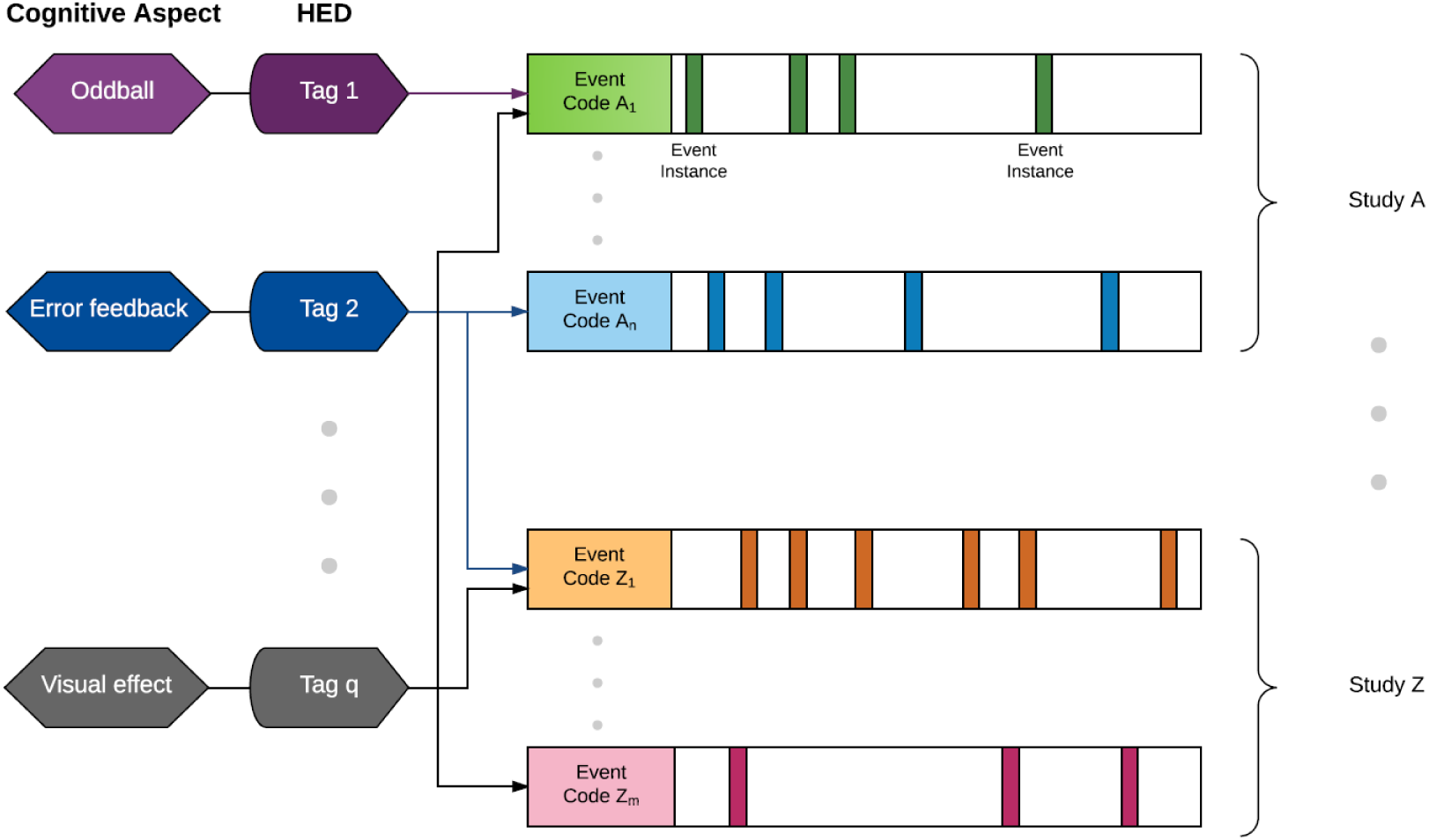
Illustration of how *HED* tags disassociate the properties of events from their study-specific codes to encode common cognitive aspects across studies.

To obtain a more detailed systematic understanding of the underlying relationships, we also implemented a second-level linear regression model (General Regression of Aspects and Details or *GRAND*) as illustrated in Figure 3. *GRAND* uses three types of factors in its design matrix: recording identifiers, study-specific event codes, and cognitive aspects (*HED* tags). Each event-related pattern has a unique study-specific event code and recording identifier, but can be associated with many cognitive aspects. These cognitive aspects (e.g., *Sensory presentation/Visual*, or *Participant/Effect/Cognitive/Target* or *Action/Button press*) are captured by the *HED* tags assigned to the event code. The event code factors capture variability due to paradigm differences across studies, while the recording factors capture subject/session/headset variability. *HED* tag factors capture commonalities across events from different studies.

**Figure 3.**
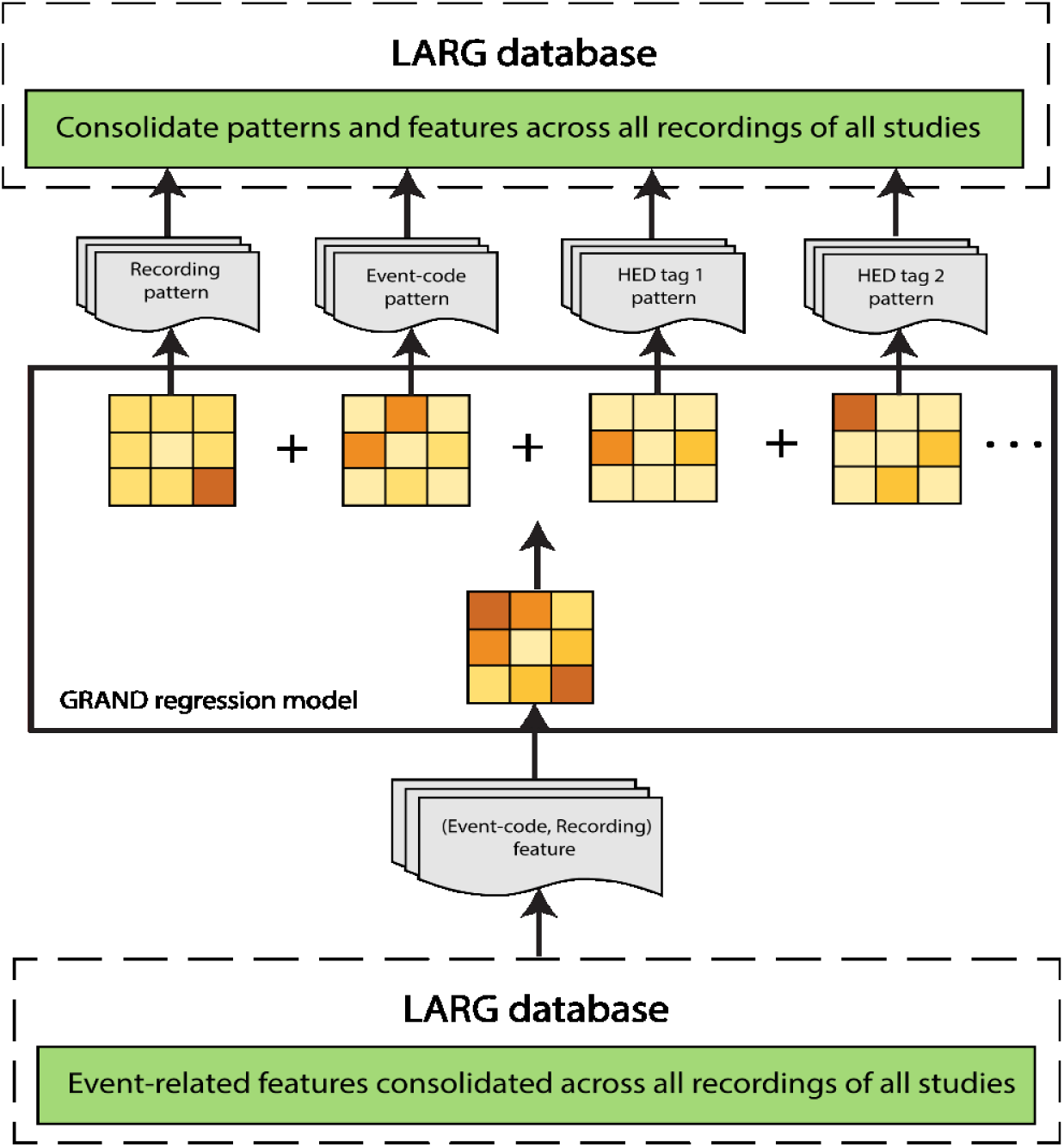
*GRAND* models each (Event-code, Recording) EEG pattern as a sum of a recording-specific pattern, a study-specific pattern associated with each unique event code in the study, and a pattern for each *HED* tag present in the annotation of the events of that recording. The patterns are added to the *LARG* database for analysis.

First level regression or averaging over recording events of a given event type produces a pattern (i.e., rERP, rERSP, ERP, or ERSP), *P*_*n*_, for each of the *N* unique (event code, recording) pairs in the data collection. Second level regression (*GRAND)* employs *M = T+ R + H* factors, where *T* is the total number of distinct study-specific event codes in the collection, *R* is the total number of recordings, and *H* is the total number of selected cognitive aspect factors, conveyed by *HED* tags. GRAND calculates the patterns, *Q*_*m*_, associated with the *M* factors based on the linear model:

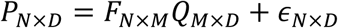

*D* is the size of the event-related pattern, and *F* is the design matrix for the regression. The optimization problem for *GRAND* is:

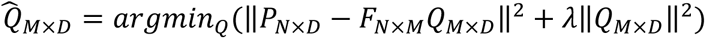

where *λ* is a regularization parameter estimated using the generalized cross validation method described previously.

Outlier (Event-code, Recording) pairs are detected and excluded from regression by an iterative method similar to that used by the first-level THOR models. On each iteration, *GRAND* computes a linear model based on non-outlier data and applies the model to *all* of the data to obtain residuals. Outliers for the iteration are defined as (Event-code, Recording) pairs with residuals exceeding a robust *z*-score of 5. A maximum of 100 iterations are performed, with an early termination criterion of no change in outlier designations after one iteration.

### 2.5 Statistical and visual analysis of event-related patterns across studies

In addition to the traditional visualizations of ERP and ERSP patterns, we used both visual and statistical analysis to explore the relationships between patterns and cognitive aspects across studies. For visualization, we applied *t-SNE* (t-distributed stochastic neighbor embedding) to project high-dimensional feature vectors onto a two-dimensional (2D) space. The idea is that given a normalization and distance metric, high-dimensional vectors that are similar to each other (e.g., relatively “near” each other in the high-dimensional space) will project to nearby points in the low-dimensional 2D subspace. A cluster in the 2D subspace thereby indicates a group of feature vectors that are more similar to each other, relative to other features.

For example, to analyze rERSPs, we calculated an rERSP for each study-specific event code and each of 26 common channels in each recording. We formed a vector by concatenating the rERSPs of these channels for a recording and normalized using *z*-scoring. The *t*-*SNE* visualizations project each normalized vector to a point in a 2D space. We color-code each point based on its metadata to identify clusters. The *t*-*SNE* visualizations of this paper use the algorithm and source code of van der Maaten and Hinton (Van Der Maaten and Hinton, 2008) (Van Der Maaten, 2014). For ERSP patterns we used logarithmically sampled frequencies and a time interval of [-1, 1] seconds relative to event onset.

We used representational similarity analysis (*RSA*) (Kriegeskorte et al., 2008) to quantify the relationship between event-related signal patterns and the *HED* tags associated with these event codes (e.g., are ERPs associated with two event codes that both contain the *HED* tag “Target” more likely to be similar than would be expected from a random association). We constructed two types of similarity matrices: one for EEG signal patterns and one for the presence of *HED* tags as illustrated in Figure 4.

**Figure 4.**
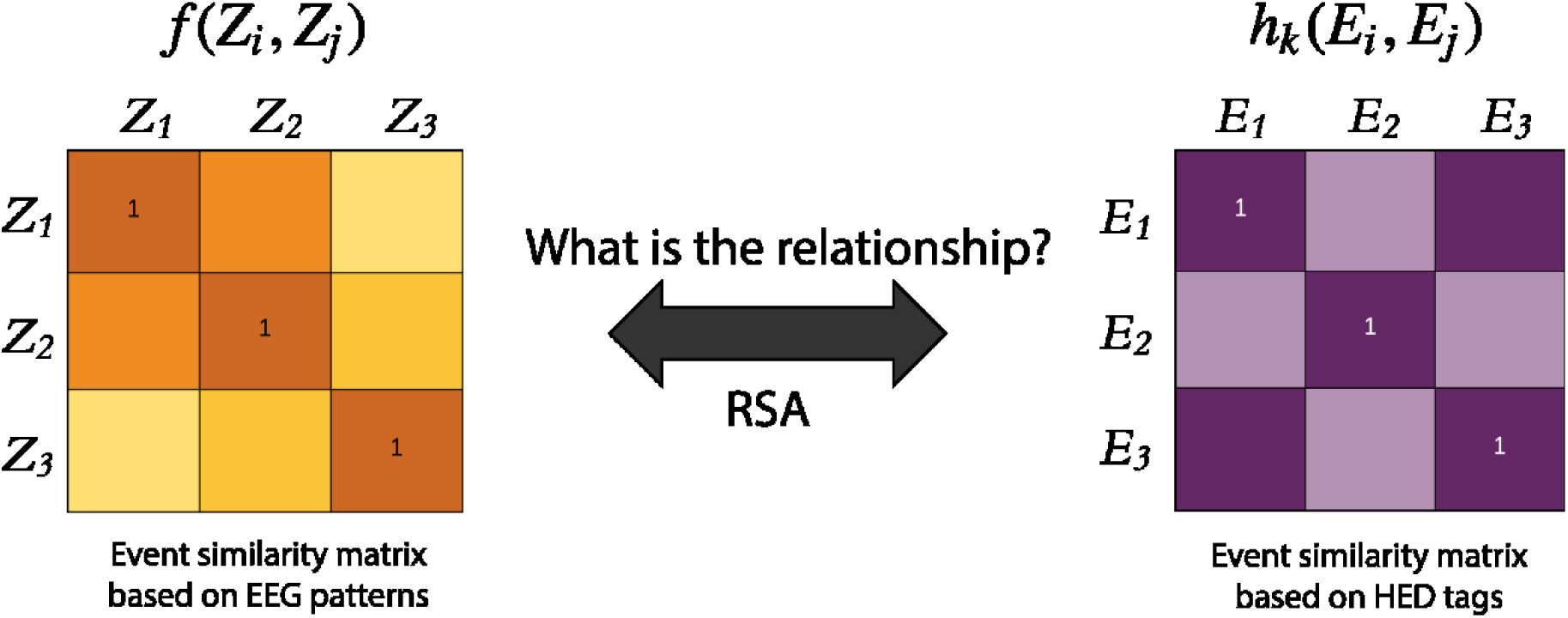
Schematic of representational similarity analysis (*RSA*) strategy used to ascertain the statistical similarity between EEG event-related signal features and their associated *HED* tags. Here *Z_i_* is a *z*-scored pattern (ERP, rERP, ERSP or rERSP) associated with a study-specific event code *E_i_*. The entries of *f* are the correlations between the patterns associated with the respective event codes. The entries of the matrix *h_k_* on the right measure the overlap in *HED* tags associated with the respective event codes, *E_i_* and *E_j_*.

Let *Z*_*i*_ be a *z*-scored feature associated with a study-specific event code, *E*_*i*_. To obtain *Z*_*i*_ for an rERP pattern, for example, we first performed a normalization using a *z*-score transform on the vectorized pattern associated with each (event type, recording) pair. We concatenated data from all channels and times before this transformation. We then averaged these *z*-score transformed rERPs for each *E*_*i*_ across all recordings in the study to obtain *Z*_*i*_ (see Figure 4, left). We computed the pattern similarity matrix *f*(*Z*_*i*_, *Z*_*j*_) as the Pearson correlations of (*Z*_*i*_,*Z*_*j*_) pairs. We computed rERSP patterns similarly by concatenating data for all channels, times, and frequencies prior to *z*- score normalization. While the similarity computation can use either trial average features (ERPs or ERSPs) or temporally regressed features (rERPs or rERSPs), regression reduces confounds due to event overlap.

The second similarity matrix represents concurrence of the presence or absence of the association of a *HED* tag *k* for event codes *E*_*i*_ and *E*_*j*_ (see Figure 4, right). Let *H*(*E*_*i*_) be the set of HED tags associated with the study-specific event code *E*_*i*_. The aspect similarity matrix, *h*_*k*_(*E*_*i*_, *E*_*j*_) is defined as:

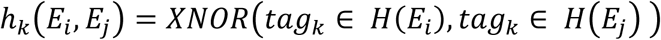

We computed *RSA* separately for each *HED* tag or *HED* tag prefix, *tag*_*k*_. Since we examined several such tags, there are a number of secondary *RSA* similarity matrices, each associated with a different tag. We first selected a set of *HED* tags based on information content and on the number of occurrences of the tag in the corpus. We then computed the secondary similarity matrix value *h*_*k*_(*E*_*i*_, *E*_*j*_) for the *k*^*th*^ eligible tag as 1 if the *k*^*th*^ tag appears in both *E*_*i*_ and *E*_*j*_ or 0 if the *k*_*th*_ tag does not appear in both *E*_*i*_ and *E*_*j*_.

We converted each similarity matrix (based on patterns or tags) to a dissimilarity matrix by performing an elementwise *1 - x* operation. Following the *RSA* steps prescribed in (Kriegeskorte et al., 2008), we determined whether there is a statistically significant relationship between dissimilarities in EEG patterns and differences in event type tag memberships. We generated 10,000 random permutations of each EEG pattern dissimilarity matrix and used Spearman correlation with each tag membership dissimilarity matrix (upper triangular part only) for the *RSA* computation. This resulted in a single significance value for each *tag*_*k*_ for each type of EEG pattern (e.g., rERP, rERSP, or the average of their two associated dissimilarity matrices).

## 3. Results

### 3.1 Temporal overlap regression reduces confounds of overlapping events

Temporal overlap regression provides an important analysis tool for reducing confounds among overlapping events in order to discover generalizable, statistically significant patterns associated with events and with cognitive aspects. Figure 5 illustrates the confound issue for a single subject in the RSVPU study for data before eye activity removal. The left column shows patterns obtained by averaging epochs associated with distinct study-specific event codes (*Target, Non-Target*, and *Eye blink/Max*), while the right column shows regressed ERPs associated with the same event codes. In these graphs and in all subsequent pattern-related graphs, gray areas indicate lack of statistical significance (*p* > 0.01, *FDR* corrected).

**Figure 5.**
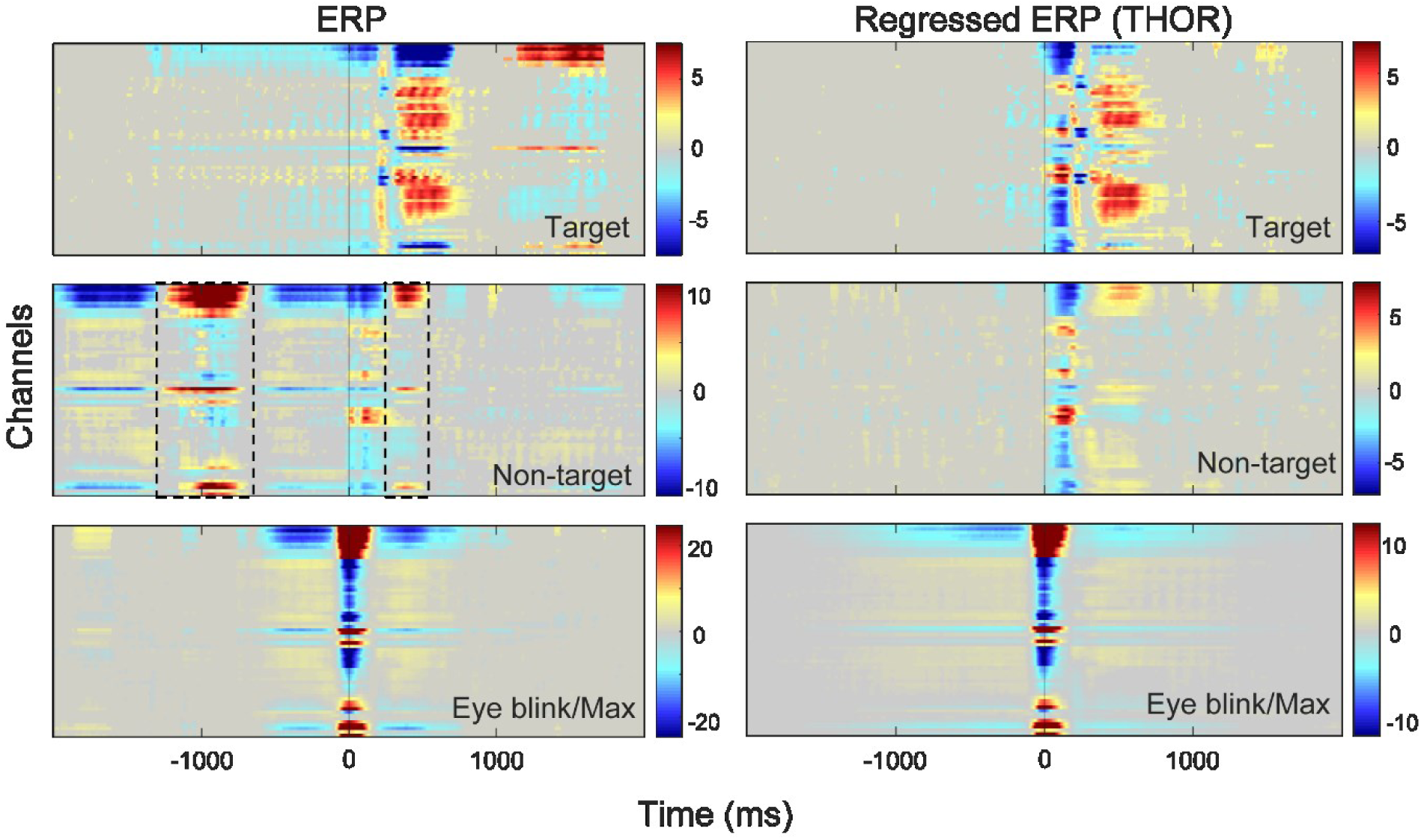
Comparison of averaging and temporal overlap regression for subject 1 in UCSD-RSVPU before removal of eye activity. The left column shows ERPs obtained by epoch averaging for events codes *Target, Non-target*, and *Eye blink/Max*, respectively. The right column shows the rERPs obtained with temporal overlap regression using the same event codes. Black dashed boxes highlight confounds.

RSVP studies are particularly prone to confounds due to the rapid presentation of interspersed target and non-target images. The 12 Hz presentation rate clearly induces an SSVEP (steady state visually evoked potential) or entrainment pattern throughout the four-second epoch, shown in the top and middle graphs of the left column of Figure 5. The SSVEP confound of rapid presentation has been mostly eliminated in the regressed ERPs displayed in the right column. The *Non-target* ERP pattern appears to be a summation of image presentation ERPs and time-shifted *Eye blink/Max* ERP patterns. The roles of blink alignment and visual system entrainment during RSVP have been widely studied (Kranczioch, 2017), and these effects clearly represent an important factor in many visually related tasks. Since our eye-removal techniques explicitly remove blinks from the time signal using regression, these confounds are less apparent in the ERPs calculated from the signal after our eye-artifact removal process.

Temporal overlap regression also removes other types of confounds and reduces spurious experimental effects as illustrated by Figure 6. These ERSPs come from channel Fz of a single recording in the DAS experiment after eye activity removal. The DAS distracted driving experiment involved the presentation of animal words on the left or right side of the dashboard requiring an immediate response (button press) for animals (targets). Because of the protocol design, a right-side presentation was likely to be surrounded by left-side presentations and vice versa. All word presentations were exactly 1500 ms apart. Hence simple averaging conflates the effects of adjacent left-side and right-side presentation (left-side and right-side presentations had similar ERSP patterns). Regression removes this confound and provides a much cleaner estimate of the underlying pattern.

**Figure 6.**
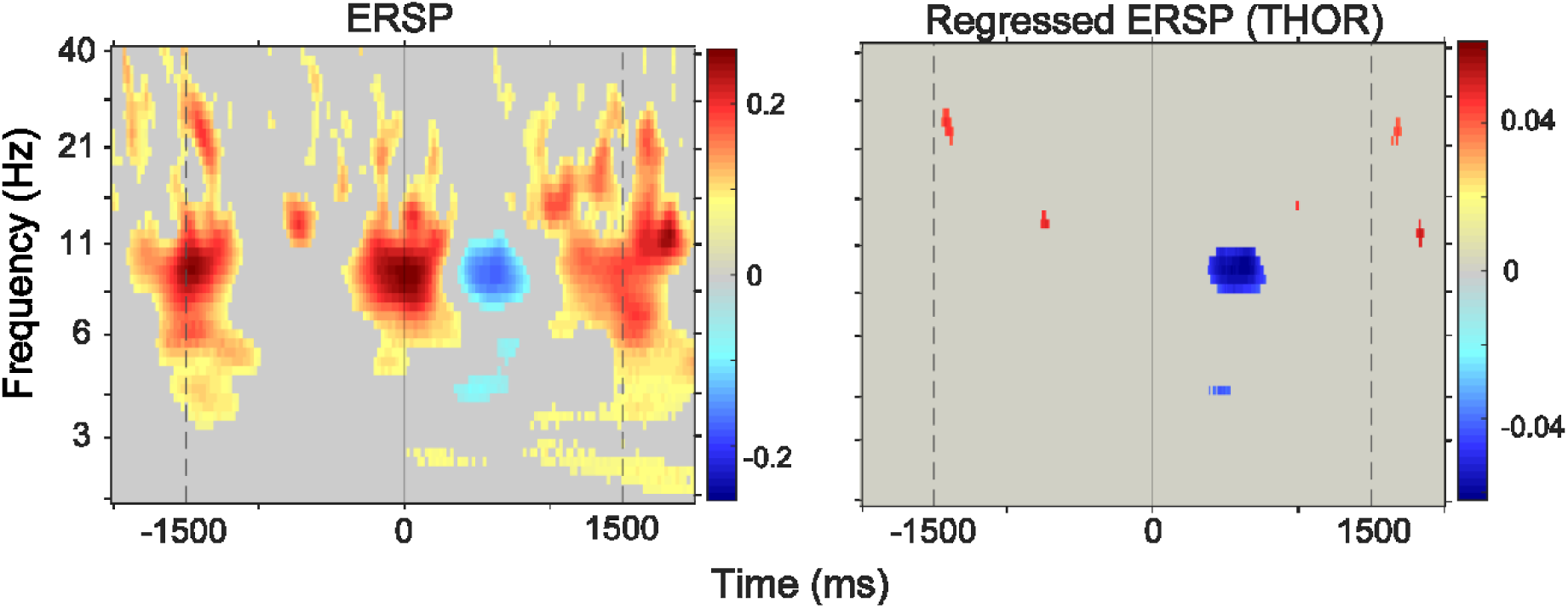
An example of a recording’s ERSP and rERSP for channel Fz time-locked to right-side presentations of visual words in the DAS experiment for the first recording in the collection. The left column shows an ERSP obtained by averaging, while the right column shows the corresponding rERSP obtained by temporal overlap regression. The features are time-locked to NCTU-DAS event code number 22.

### 3.2 Regressed cognitive aspects show similar neighborhood structure across studies

The previous section illustrated how temporal overlap regression exposes and reduces experimental confounds in both ERP and ERSP analysis for single recordings. To understand how cognitive aspects isolate similar EEG patterns across recordings and studies, we used combined temporal overlap regression and *t-SNE* visualization (Figure 7) to project ERSP features from the entire collection onto a two-dimensional subspace and examined whether patterns labeled with the same cognitive aspects projected to similar locations.

**Figure 7.**
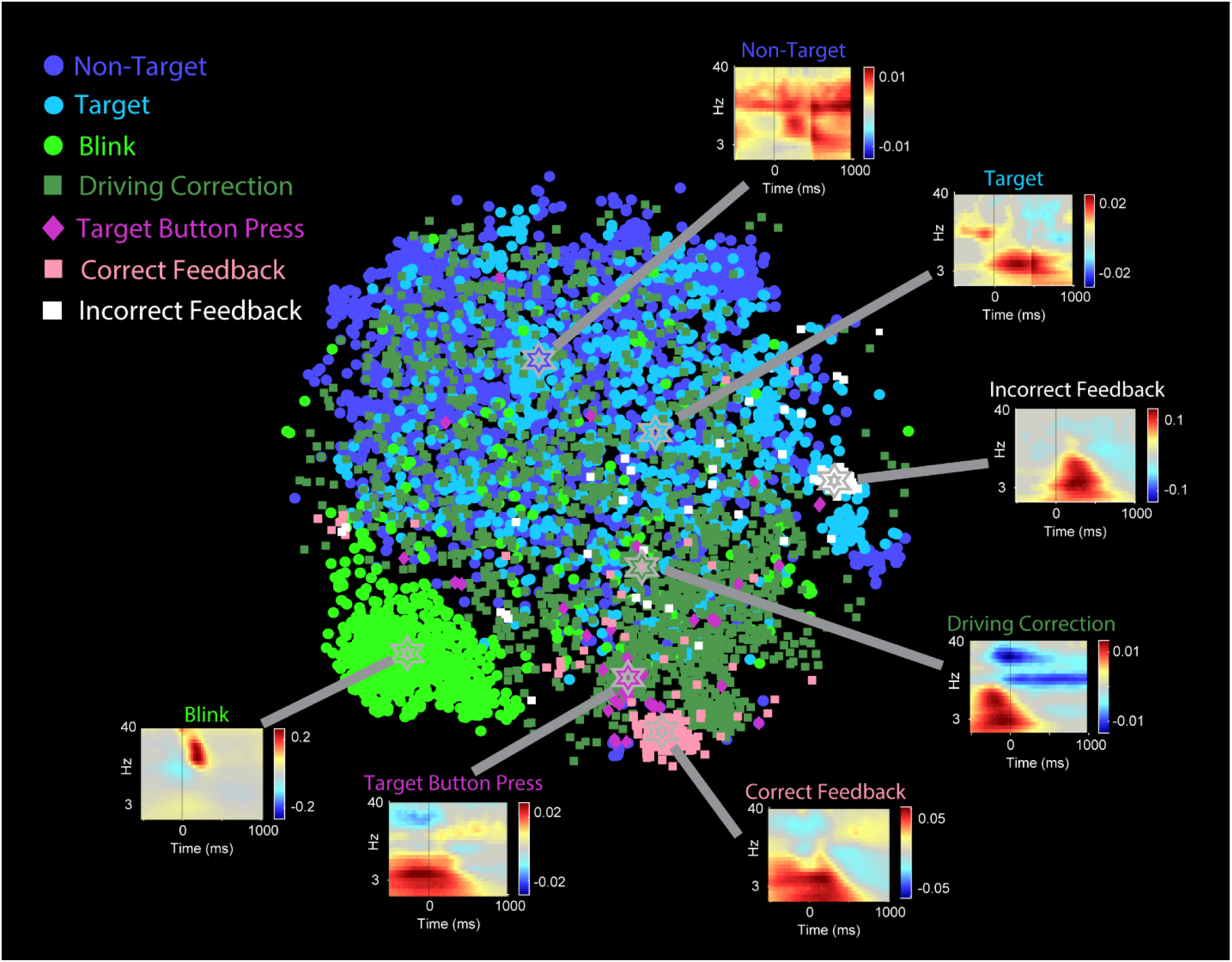
2D *t*-*SNE* visualization of rERSPs across all 17 studies for the 26 common channels. Each point represents a vector formed by the rERSPs concatenated over all channels and frequencies for the interval [-1, 1] seconds corresponding to a study-specific event code for one recording. Points are color-coded by the most prominent *HED* tag associated with the event code. The median coordinates of points associated with a particular HED tag are marked with a star. Inset plots of the average of the rERSPs of all points associated each HED tag are shown for channel FCz.

The *t-SNE* projection of Figure 7 represents a dramatic dimensionality reduction from the original 26×50×40×2 = 104,000 dimensional space down to 2 dimensions. However, the strong clustering associations of these projections with *HED* tags suggest significant pattern similarity for the corresponding rERSPs. Points of the same color that are tightly clustered and whose median positions fall close to the visual centers of the respective clusters have few outliers, suggesting that the corresponding rERSP features have very similar behavior across subjects and studies. Points of the same color that are more widely spread and whose medians do not fall within one of the prominent sub-clusters, likely reflect the diverse nature of the experimental paradigms that share the associated cognitive aspect.

The most prominent overall cluster, the green *Blink* cluster, includes points associated with every study. Blink event codes were inserted automatically into all studies by *BLINKER* and labeled with the *HED* tag *Action*/*Eye blink/Max*. Even though our pipeline aggressively removes eye-activity in the time-domain using *EyeCatch* and *BLINKER*, frequency-domain features still contain prominent and consistent blink-locked patterns. These patterns may originate not only from the eye, but also from the cortex in anticipation of or in response to the blink, as reported in the fMRI literature (Berman et al., 2012; Hupé et al., 2012; Peter et al., 2010). The existence of these robust rERSP patterns after time-domain eye-activity suppression highlights the need for temporal regression in conjunction with blink detection when computing ERSPs. Otherwise, the non-uniform distribution of blinks (i.e., more or less likely after particular event types), could lead to significant cofounds and misleading ERSP patterns. Task induced eye movements such as saccades were not regressed out in this analysis and may also lead to confounds. However, as many of the events in this data-collection come from fixation-constrained studies, blinks may be a more systematic confound.

The pink cluster, labeled *Correct Feedback* in Figure 7, corresponds to the *HED* tag *Participant/Effect/Cognitive/Feedback/Correct*. The points in this cluster are primarily from NCTU-DAS and NCTU-DD studies. Interestingly, this pink cluster and the dark green cluster labeled *Driving Correction* (*HED* tag *Action/Control vehicle/Drive/Correct*) are relatively close in this visualization, as is the magenta cluster *Button Press* (*HED* tags *Action/Button press* or *Action/Button hold*).

The distribution of *Participant/Effect/Cognitive/Target* and *Participant/Effect/Cognitive/Non-target HED* tags (labeled *Target* and *Non-target*, respectively) is also interesting. Most of the 17 studies have these tags in at least one event code, and their pervasiveness is reflected in the cyan and dark blue points spread throughout Figure 7. Interestingly, the averaged rERSPs corresponding to these HED tags each show distinct discontinuities at 500 ms.

Figure 8 examines the source of this discontinuity in more detail. The left column of Figure 8 shows the results of averaging the ERSPs computed over the recordings in four different studies. The right column shows the results of performing the same averages for regressed ERSPs. The rERSP average for NCTU-DAS shows a marked discontinuity at 500 ms, while the other three studies do not. The experimental protocol for NCTU-DAS was such that non-targets were always presented exactly 1500 ms before and after a target. The interaction of this confound with the -2 to 2 second regression intervals is the cause of this discontinuity (other regression intervals will cause similar of discontinuities).

**Figure 8.**
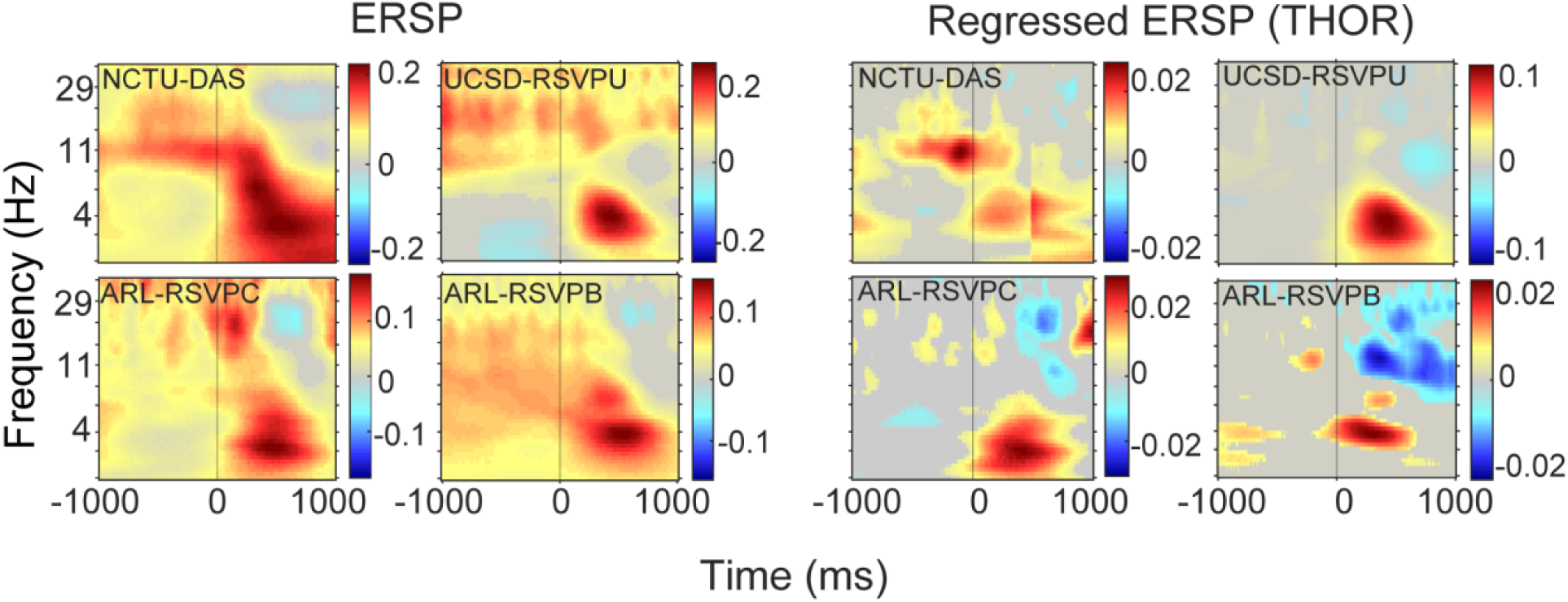
Representative study averages of ordinary ERSPs (left group) and rERSPs (right group) for channel FCz for the *Participant/Effect/Cognitive/Target* cognitive aspect. The studies in each group are NCTU-DAS, UCSD-RSVPU, ARL_RSVPB, and ARL-RSVPC (clockwise starting from the upper left corner).

Interestingly, the averaged ERSP for NCTU-DAS does not show the discontinuity. However, the ERSP appears to have a more complex and extended structure (with nearly every pixel being significant), indicating potential confounds.

### 3.3 Regression of rERPs and rERSPs extracts common cognitive aspects across studies

This section compares the effectiveness of aspect averaging and temporal overlap regression for extracting common features of cognitive aspects across studies. We apply a two-level approach: modeling patterns at the recording level and then combining patterns across all recordings in all studies.

The ERSP patterns at the first level (corresponding to study-specific event codes and individual recording) may be computed by either averaging or temporal regression. ERSP patterns associated with cognitive aspects at the second-level (combination of patterns across studies) may be computed either by averaging or regression. Aspect averaging collects and averages patterns whose study-specific event code is annotated by a particular *HED* tag. Second level regression uses the GRAND model, described in the Methods section, to account for experimental protocol, recording details, and cognitive aspects in the design matrix. These results do not include NCTU-DAS because of the discontinuity issue described above.

Figure 9 shows a cross-study comparison for the *Participant/Effect/Cognitive/Target* cognitive aspect on channel Fz. Focusing on the results of second-level modeling of rERSPs (lower graphs), we observe a response with a region of increased spectral power for 2 to 6 Hz over the 200 to 500 ms time range. This is similar to a dramatic increase in frontal delta (2 to 4 Hz) over the 100 to 500 ms time range for the active oddball task reported by Cahn et al. (Cahn et al., 2013). Cahn et al. also report a strong suppression of late low-alpha (8 to 10 Hz, 500 to 900 ms) similar to observed in Figure 9. The beta suppression (15 to 30 Hz, 300 to 1000 ms) was observed by Güntekin et al. in healthy patients, but not in those with mild cognitive impairment (Güntekin et al., 2013). Touryan et al. observed a similar pattern of frontal beta suppression in their calculations of FRPs (fixation related potentials) in a guided detection task (Touryan et al., 2017).

**Figure 9.**
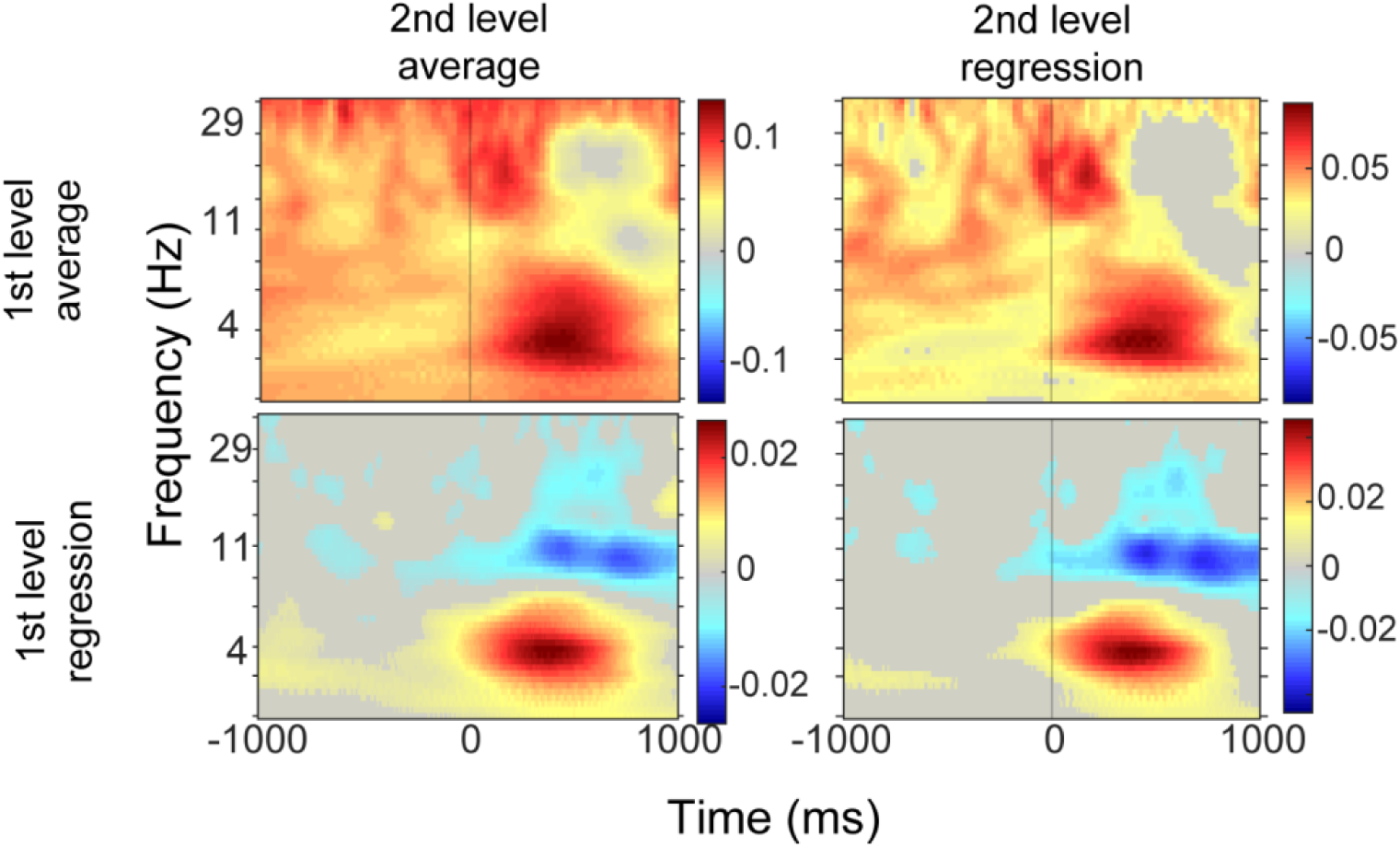
Comparison of four methods for combining recording-level ERSPs across studies. All graphs are based on the *Participant/Effect/Cognitive/Target* cognitive aspect for channel Fz across all studies except NCTU-DAS. The top row uses recording-level ERSPs obtained by aspect averaging, while the bottom row uses recording-level rERSPs. In each case the left column averages all ERSPs or rERSPs, while the right column regresses ERSPs or rERSPs.

Significantly increased frontal delta post-stimulus power related to target detection have been observed in healthy subjects for both visual and auditory targets (Caravaglios et al., 2008) (Güntekin and Başar, 2016). A difficulty in making a direct comparison with those studies and the present work is that many of them treat the pre-stimulus signal as the baseline and remove the average from the signal. We treat the pre-stimulus and post-stimulus periods symmetrically since our method uses a baseline amplitude computed over the whole recording. Notice that the same basic features occur regardless of whether aspect averaging or GRAND regression is used at the second level for rERSP features (bottom row). However, it appears from Figure 9 that confounds introduced at the first level by simple averaging at the recording level (top row of Figure 9) wash out relevant cross-study comparisons when these features are combined.

Figure 10 compares various second-level modeling approaches for cognitive aspect *Action/Control vehicle/Drive/Correct* on channel Fz. This *HED* tag annotates the point at which a driver begins to correct for a lane deviation perturbation in a driving simulation.

**Figure 10.**
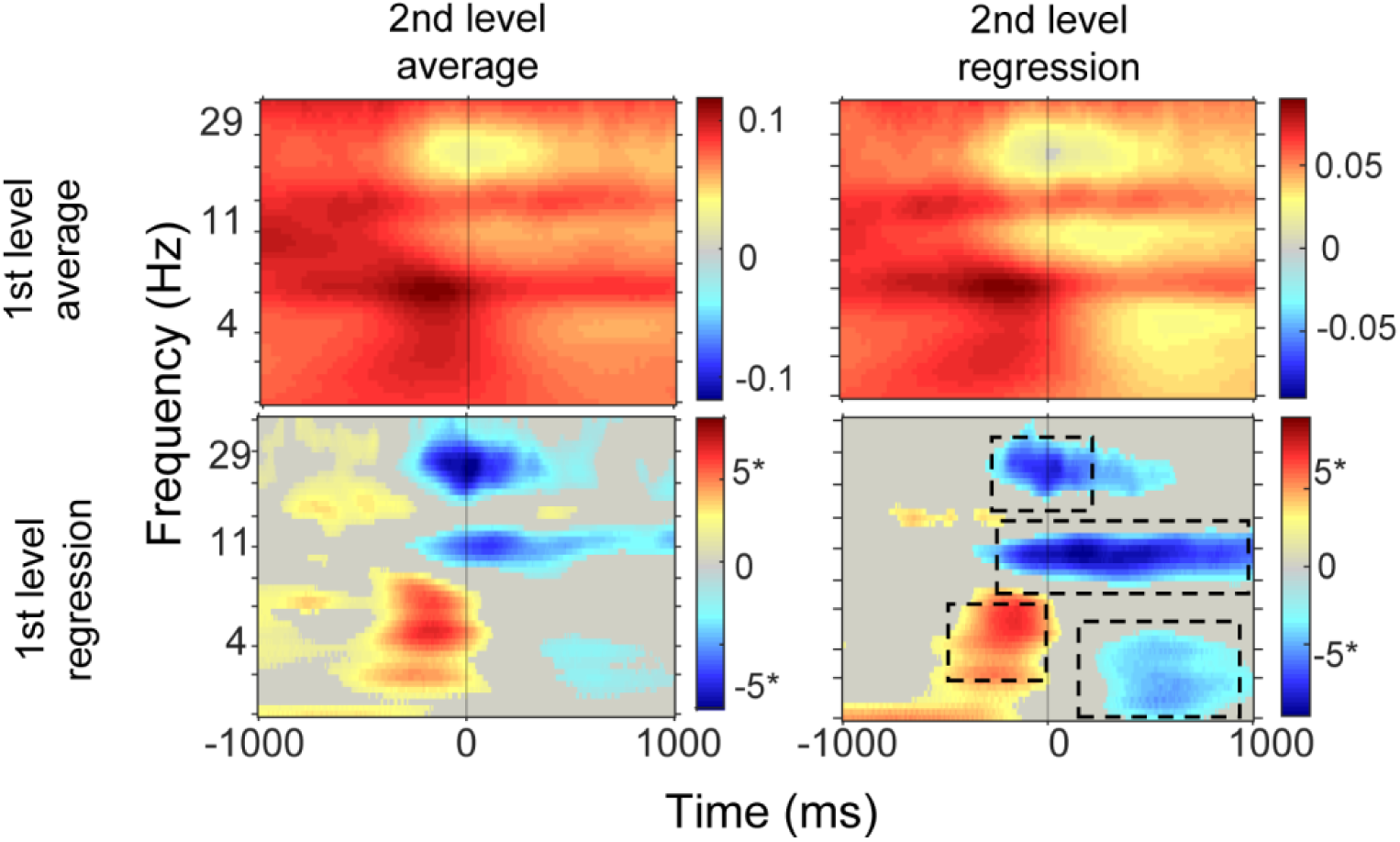
Comparison of four methods for combining recording-level ERSPs for the *Action/Control vehicle/Drive/Correct* cognitive aspect on channel Fz across all studies except NCTU-DAS. Top row uses recording-level ERSPs obtained by aspect averaging, while the bottom row uses recording-level rERSPs. In each case, the left column averages ERSPs or rERSPs, while the right column regresses ERSPs or rERSPs.

Lin et al. observed an increase in frontal power for 0 to 15 Hz in the 0.5 seconds prior to starting the correction (Lin et al., 2011). This observation is consistent with the results of Figure 10 obtained using rERSP features (bottom row of Figure 10). The cross-study responses also show strong beta frontal suppression time-locked to the correction as well as a sustained frontal high-alpha suppression during the actual accomplishment of the correction. Lin et al. also observed frontal suppression of frequencies in the range 8 to 30 Hz from 1000 ms to 2000 ms after the driver starts the correction. This period is outside the range investigated in Figure 10. These features appear in both aspect averaging and second-level regression of rERSPs, but are more clearly defined using second-level regression. Cross-study combination of ERSP features in both Figures 9 and 10 exhibit echoes of the rERSP cross-study phenomena sitting on top of a baseline response level.

Figure 11 compares the results for cognitive aspect *Action/Button press* or *Action/Button hold* for channel C3. These results are consistent with results reported by Makeig et al. (Makeig et al., 2004) and Delorme et al. (Delorme et al., 2007) showing a large delta/theta power increase preceding the button press.

**Figure 11.**
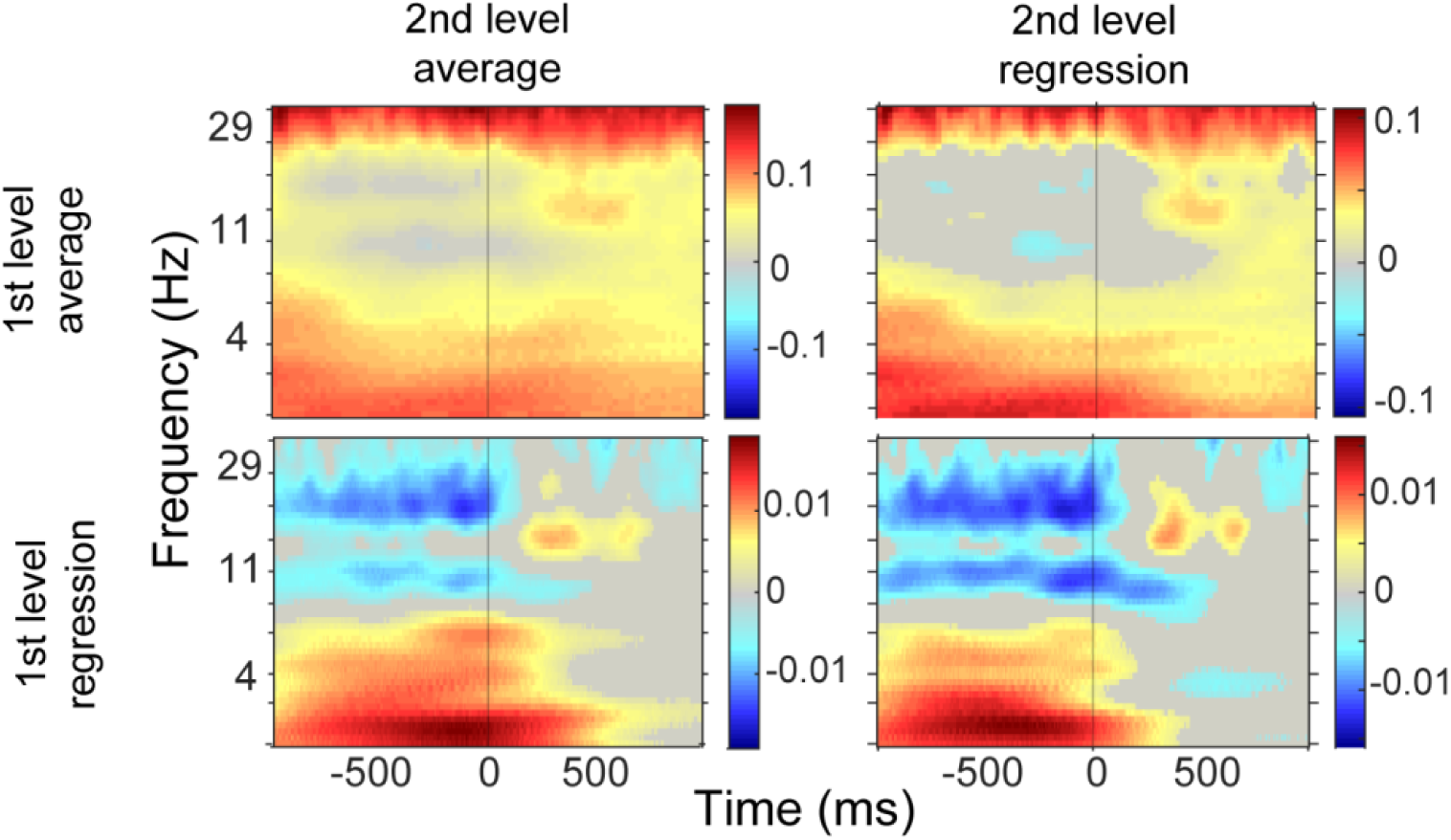
Comparison of four methods for combining recording-level ERSPs across studies. All graphs are based on the *Action/Button press* cognitive aspect for channel C3 across all studies except NCTU-DAS. The top row uses recording-level ERSPs obtained by aspect averaging, while the bottom row uses recording-level rERSPs. In each case the left column averages all ERSPs or rERSPs, while the right column regresses ERSPs or rERSPs using *GRAND*.

Makeig et al. also report that the overall power spectrum of the channel grand average shows suppression in the range 15 to 30 Hz at the button press and suppression in the 8 to 15 Hz range slightly after the button press. The cross-study spectra derived from rERSPs (lower row of Figure 11) also show this general behavior, although here it appears before the button press. The timing and the spread of these spectral phenomena may be due to variations in the reporting times of when the button is considered pushed in the various studies. Another spectral feature, corresponding to increased spectral power for 8 to 29 Hz in the range 400 ms to 1000 ms after the button press appears in all of the approaches of Figure 11, as well is in the grand channel averages reported by Makeig et al.

### 3.4 *RSA* shows statistically significant relationships between cognitive aspects and EEG patterns across studies

As summarized in Table 1, the 17 studies analyzed in this paper contained 1,155 recordings with 7,770,851 event instances. The collection contained 664 distinct study-specific event codes annotated by 192 unique *HED* tags, excluding the required or recommended *HED* informational annotations (*Event/Description, Event/Label, and Event/Long name*). Before exploring or quantifying the event-related features of each cognitive aspect, we used *RSA* to provide a global assessment and validation of the ontology. *RSA* provides a statistical method of assessing the correspondence between EEG patterns and cognitive aspects without making assumptions about the specific features (Table 2). Significant correspondences tend to be concentrated in the *Action, Participant/Effect*, and *Visual* categories.

**Table 2.** *RSA* significance of relationship between selected *HED* tags and EEG patterns across 17 studies. Tags appear in decreasing order of information content as measured by Shannon entropy.

| Tag | Total number | Codes | Studies | Entropy | rERP + rERSP | rERP | rERSP |
| --- | --- | --- | --- | --- | --- | --- | --- |
| *Item* | 1246K | 296 | 17 | 4.76 | 0.0001* | 0.0001* | 0.0001* |
| *Event/Category/Experimental stimulus* | 775K | 218 | 17 | 4.28 | 0.0001* | 0.0001* | 0.0001* |
| *Item/Object/Vehicle/Car* | 629K | 131 | 10 | 4.22 | 0.0064* | 0.0001* | 0.0777 |
| *Participant/Effect* | 792K | 210 | 14 | 4.18 | 0.0037* | 0.0001* | 0.0328 |
| *Action* | 1220K | 183 | 17 | 4.14 | 0.0001* | 0.0033* | 0.0001* |
| *Sensory presentation* | 605K | 166 | 17 | 3.88 | 0.0001* | 0.0001* | 0.0009* |
| *Attribute/Object control* | 242K | 89 | 10 | 3.83 | 0.8697 | 0.0134 | 0.9871 |
| *Action/Control vehicle* | 457K | 92 | 10 | 3.79 | 0.078 | 0.5021 | 0.0289 |
| *Action/Control vehicle/Drive* | 439K | 88 | 10 | 3.71 | 0.0784 | 0.5686 | 0.0233 |
| *Event/Category/Participant response* | 451K | 95 | 17 | 3.68 | 0.0926 | 0.64 | 0.038 |
| *Sensory presentation/Visual* | 483K | 130 | 15 | 3.63 | 0.0001* | 0.0001* | 0.0114* |
| *Attribute/Object control/Perturb* | 211K | 63 | 9 | 3.58 | 0.8717 | 0.0069* | 0.9903 |
| *Sensory presentation/Visual/Rendering type/Screen* | 457K | 108 | 9 | 3.46 | 0.0013* | 0.0001* | 0.051 |
| *Participant/Effect/Visual* | 445K | 104 | 9 | 3.40 | 0.0006* | 0.0001* | 0.072 |
| *Participant/Effect/Cognitive/Target* | 293K | 78 | 9 | 3.28 | 0.9788 | 0.5257 | 0.981 |
| *Action/Button press* | 223K | 63 | 11 | 3.27 | 0.8713 | 0.9246 | 0.6664 |
| *Participant/Effect/Body part/Arm/Hand* | 209K | 58 | 10 | 3.14 | 0.6295 | 0.8723 | 0.4559 |
| *Participant/Effect/Cognitive/Oddball* | 272K | 42 | 6 | 3.03 | 0.994 | 0.609 | 0.9949 |
| *Participant/Effect/Body part/Arm/Hand/Finger* | 190K | 54 | 10 | 2.98 | 0.6693 | 0.9606 | 0.4254 |
| *Participant/Effect/Cognitive/Oddball/ Button pressing for target* | 262K | 37 | 4 | 2.92 | 0.9739 | 0.3046 | 0.9856 |
| *Action/Button press/Keyboard* | 185K | 27 | 7 | 2.87 | 0.9314 | 0.9781 | 0.843 |
| *Participant/Effect/Cognitive/Non-target* | 356K | 46 | 8 | 2.86 | 0.007 | 0.404 | 0.0063 |

A \* indicates an association between the respective EEG pattern(s) and cognitive aspects (*HED* tags) that is significantly stronger than random ( $p < 0.01$ ) after *FDR* multiple comparison correction across all $p$ values in the table.
Statistically significant associations that occur ( $p < 0.01$ ) only when correcting for multiple comparison using $p$ values of the same pattern type are shown in black. Light gray type indicates non-significance.

The *Action* tags tend to be associated with participant responses, while *Participant/Effect* tags tend to be associated with stimulus type events. These results are consistent with the types of studies analyzed for this paper: many of the studies have a visual target component, a large subset of studies involve driving in a simulator with lateral vehicle perturbations, and many experiments require some type of button or key press as an explicit subject response. Several of these tags appear in only a few event codes and hence have a lower “diversity”, increasing their susceptibility to the peculiarities of specific event types.

## 4 Discussion/conclusion

The results of this paper were based on 17 studies from 6 sites at 4 different institutions that captured approximately 7.8 million event instances from several different experimental paradigms. This work demonstrates the viability of event-related mega-analysis for EEG and presents a variety of applicable methods. The core analysis of this paper explores the relationship of cognitive aspects as represented by *HED* tags to EEG event-related patterns (ERPs, rERPs, ERSPs, and rERSPs) across a large corpus. At the first (recording) level, we compute these patterns separately for each study-specific event code and each recording. As observed by other authors (Burns et al., 2013) (Ehinger and Dimigen, 2018), temporal overlap regression provides much cleaner event-related patterns and substantially reduces confounding effects from nearby events.

We constructed a database of these patterns, labeled by (Event-code, Recording) tuples and used this database for analysis (*RSA*) and visualization (*t*-*SNE*). Application of *RSA* analysis to rERPs and rERSPs reveals statistically significant relationships between the presence of common *HED* tags and the structure of the associated EEG event-related signal patterns. These relationships are exhibited in more detail through the ERSPs.

We combined patterns at a higher (multi-study) level by regressing the resulting first-level patterns onto *HED* tag, study-specific event codes, and recording factors to create a two-level hierarchical model of the collection as a whole (*GRAND*). The recording factors encapsulate both subject and session information. The study-specific event-code factors encapsulate the effects of study-specific differences on brain dynamics, associated with common cognitive aspects, caused by differences in paradigms and the protocols implementing these paradigms. Since each study used the same headset at a given site, the study-specific event-codes also capture common headset information. These factors allow the assessment of the variability of patterns across studies having the same paradigm (but different protocols or equipment), enabling a quantitative assessment of generalizability. Hierarchical general modeling has been successfully used to test for statistically significant ERP and ERSP patterns across subjects (Pernet et al., 2011), but the introduction of *HED* tags as cognitive aspects in the regression models allows decoupling of study-specific paradigm differences from fundamental or universal responses.

This work demonstrates the importance of using temporal overlap regression at the recording level when comparing event-locked features across studies, within studies, or even at the single recording level. As demonstrated in Figure 5, blinks are ubiquitous confounds that are not always randomly distributed with respect to other stimuli, particularly in visually intensive tasks. We believe that blinks should be treated as events in their own right and regressed out as part of the processing. Further, overlap regression can reveal unexpected features resulting from protocol design that are obscured by simple averaging (Figure 6 and Figure 8).

As demonstrated in Figures 9 through 11, the consequence of not taking into account confounds at the recording level propagate downstream in the form of diffuse structure when recording-level results are combined in mega-analysis. Thus, clean results at the recording level are essential for downstream comparisons.

The particular organization of the pattern database at the first level by (Event-code, Recording) allows new studies to be easily incorporated and first-level features computed in a scalable manner. Our database effectively consists of a map that associates identifying characteristics (event codes and recordings) with various EEG features. The second-level regression over the database is, therefore, a relatively low-cost operation compared with preprocessing and first-level pattern processing. This is particularly important as the 17 studies included in this work are heavily oriented towards visual processing, target detection, and driving. As a broader selection of well-annotated raw EEG becomes available, we hope to analyze additional patterns in the context of the collection. Pattern matching across ERP datasets has been proposed by Liu et al. (Liu et al., 2012). However, we believe that because of the variability and confounding factors of EEG as a neuroimaging modality, such matching cannot be effective without a common processing pipeline and consistently implemented temporal overlap regression.

The association of study-specific event-codes with cognitive aspects represented by *HED* tags is a crucial step in this cross-study analysis and, once completed, allows fully-automated downstream processing. However, successful analysis is predicated on the assumption that *HED* annotation is done in a consistent manner. The current 17 studies were annotated by one or two people and checked by a small group for consistency. In our experience, annotating a study that uses an unfamiliar paradigm requires thought and discussion, but once reviewed, annotation of other studies using variations of the paradigm is relatively easy.

*HED* has recently been incorporated into the BIDS standard (Gorgolewski et al., 2016) as the method for annotating events for all neuroimaging modalities (fMRI, fNIRS, MEG, EEG). We have developed a *HED* annotation guide (*HED tags and related tools*, 2018) and continue to add examples as we gain experience annotating studies involving new experimental paradigms. We are beginning to add identifiers from the Cognitive Atlas (*Cognitive atlas*, 2018) directly into the *HED* schema to better enable more sophisticated methods of automated event processing in the future. *HED* has a variety of validation tools and we are also working to deploy web-based *HED* validation.

EEG mega-analysis is a nascent subfield and this is a work in progress. The automated nature of this approach and the assembly of a large corpus of data will permit more systematic investigation of brain dynamics associated with cognitive phenomena, along with an assessment of generalizability of results obtained from single studies or paradigms. We believe that a barrier to community adoption of preprocessing standards is the lack of benchmarking and systematic evaluation of methodological effects. This work and related analysis of continuous EEG proposed in a companion paper (Bigdely-Shamlo et al., 2018) provide a potential framework for systematically evaluating the influence and effectiveness of various preprocessing strategies.

## 5. ACKNOWLEDGMENTS

The authors would like to thank Scott Makeig for inspiring this work. We would like to acknowledge Tony Johnson and Michael Dunkel of DCS Corporation for their careful assembly and curation of the ARL data. We would also like to acknowledge Ching-Teng Lin and Jung-Tai King of NCTU and the other experimenters who contributed their data to this effort. This work received computational support from UTSA’s HPC cluster Shamu, operated by the Office of Information Technology. Research was sponsored by the Army Research Laboratory and was accomplished under Cooperative Agreement Number W911NF-10-2-0022 (CAST 076910227001). The views and the conclusions contained in this document are those of the authors and should not be interpreted as representing the official policies, either expressed or implied, of the Army Research Laboratory or the U.S Government. The U.S Government is authorized to reproduce and distribute reprints for Government purposes notwithstanding any copyright notation herein.

